# Mutation patterns in SARS-COV-2 Alpha and Beta variants indicate non-neutral evolution

**DOI:** 10.1101/2022.02.28.482283

**Authors:** Monika Kurpas, Marek Kimmel

## Abstract

Due to the emergence of new variants of the SARS-CoV-2 coronavirus, the question of how the viral genomes evolved, leading to the formation of highly infectious strains, becomes particularly important. Two early emergent strains, Alpha and Beta, characterized by a significant number of missense mutations, provide natural testing samples.

In this study we are exploring the history of each of the segregating sites present in Alpha and Beta variants of concern, to address the question whether defining mutations were accumulating gradually leading to the formation of sequence characteristic of these variants.

Our analysis exposes data features that suggest other than neutral evolution of SARS-CoV-2 genomes, leading to emergence of variants of concern. We observe only small number of possible combinations of mutations indicating rapid evolution of genomes. In addtion, mutation patterns observed in whole genome samples of Alpha and Beta variants also indicate presence of stronger selection than in remaining genome samples.

## 1 Background

Severe Acute Respiratory Syndrome Coronavirus 2 (SARS-CoV-2) causing current COVID-19 pandemic as typical RNA virus is expected to mutate at a pace of 10^-4^ nucleotide substitutions per site per year [18, 12].

Although most of these mutations are either deleterious or neutral, some of them may impact transmissibility and infectivity of the emerging strain. Accumulation of mutations may lead also to immune escape increasing likelihood of reinfection. These features are observed in several strains, called ‘variants of concern’ (VOCs) characterized by sets of mutations.

### 1.1 B.1.1.7 (Alpha) variant

B.1.1.7 variant, later recognized as a variant of concern, was first detected in November 2020 in a sample taken on September 20, 2020 in the United Kingdom. With transmissibility increased by 43-90% [4] and about twofold replicative advantage [7], Alpha variant began to spread, quickly outnumbering the original Wuhan strain. B.1.1.7 variant is characterized by 14 non-synonymous mutations and 3 deletions [8, 15] (Tab. 1).

**Table 1:** Defining mutations of B.1.1.7 (Alpha) variant.

| Gene | Nucleotide | Amino acid |
| --- | --- | --- |
| ORF1ab | C3267T | T1001I |
|  | C5388A | A1708D |
|  | T6954C | I2230T |
|  | 11288–11296 deletion | SGF 3675–3677 deletion |
| Spike | 21765–21770 deletion | HV 69–70 deletion |
|  | 21991–21993 deletion | Y144 deletion |
|  | A23063T | N501Y |
|  | C23271A | A570D |
|  | C23604A | P681H |
|  | C23709T | T716I |
|  | T24506G | S982A |
|  | G24914C | D1118H |
| ORF8 | C27972T | Q27stop |
|  | G28048T | R52I |
|  | A28111G | Y73C |
| N | 28280 GAT→CTA | D3L |
|  | C28977T | S235F |

### 1.2 B.1.351 (Beta) variant

Another of several SARS-CoV-2 variants believed to be of particular importance, was first detected in the Nelson Mandela Bay metropolitan area of the Eastern Cape province of South Africa in October 2020. The B.1.351 variant is characterized by 19 mutations, with 9 of them in the Spike protein coding region [19] (Tab. 2).

**Table 2:** Defining mutations in Spike coding region of B.1.351 (Beta) variant.

| Gene | Nucleotide | Amino acid |
| --- | --- | --- |
| Spike | C21614T | L18F |
|  | A21801C | D80A |
|  | A22206G | D215G |
|  | G22813T | K417N |
|  | G23012A | E484K |
|  | A23063T | N501Y |
|  | A23403G | D614G |
|  | C23664T | A701V |
|  | 22281-22289 | deletion of three amino acids<br>at positions 242 to 244 |

In this study we are exploring the history of each of the segregating sites present in Alpha and Beta VOCs. We are trying to answer the question whether defining mutations were accumulating gradually until they form a sequence characteristic of Alpha or Beta variant, or whether this phenomena can be explained by recombination of two genomes with subsets of mutations.

We also check whether mutation patterns observed in whole genome samples of viral variants classified as VOCs indicate the presence of stronger selection than in non-VOC samples.

## 2 Methods

### 2.1 Multiple Sequence Alignment and sample preparation

The analysis was carried out using 384,741 nucleotide sequences of SARS-CoV-2 genomes, downloaded from the GISAID (Global Initiative on Sharing Avian Influenza Data) database [17, 2]. The list of accession numbers for most important sequences can be found in Supplementary Information. Samples were dated from the period 24 December 2019 to 22 January 2021.

The sequences were aligned using MAFFT Multiple Sequence Alignment Software Version 7 [9, 10] using NC_045512.2 – the first sequenced SARS-CoV-2 genome from Wuhan [3] as a reference sequence to accelerate the calculations and to recognize gene positions inside the created Multiple Sequence Alignment (MSA).

### 2.2 Algorithms to generate weekly statistics of viral genomes

We created statistics for each week since beginning of the pandemic writing down the total number of genomes and also the number of Alpha and Beta-variant genomes in given week.

### 2.3 Studies of segregating sites

Segregating sites, characteristic of Alpha and Beta SARS-CoV-2 variants (see Background section) were identified from the alignment based on comparison with reference sequence. The length of the complete sequence in the alignment is 61,796 nucleotides. The length of the Alpha variant segregating sites subsequence is 33 nucleotides (including positions of deletions), while Beta variant segregating sites subsequence has 10 nucleotides (including positions of deletions).

We reviewed all 384,741 subsequences of SARS-CoV-2 genomes. For each position in the sub-sequence we checked whether given genome has VOC-defining mutation in corresponding place. Then, if this was the case, we saved the accession number and collection date of such genome. Having these data enabled us to quantify the change in the abundance of individual mutations over time, and to study possible combinations of 2, 3, 4 etc. mutations present together in one genome as well as to determine the dates when such combinations arose. We compared observed counts of combinations in tested samples with expected number of combinations, given the count of segregating sites.

### 2.4 Studies on the site frequency spectra

In order to check whether there is higher selection pressure among genomes belonging to the VOC strain than to the remaining strains we divided our dataset into two groups: Alpha strain genomes and remaining genomes (in the second experiment we did the same for Beta strain genomes). Then we divided both groups by weeks and chose weeks with suitable number of sequenced VOC strain genomes. Results of analysis of number of genomes sequenced in given week are presented in Fig. S1. The selection criterion for the week was the number of VOC genomes, which shouldn’t be larger than 500 (due to computational limitations) and shouldn’t exceed number of non-VOC genomes sequenced in a given week. In the case of Alpha these were weeks 45 (115 genomes), 46 (292 genomes) and 49 (392 genomes) since the beginning of the pandemic.

For each of these weeks we randomly chose equal number of non-VOC genomes, to keep both samples the same size. For each sample we calculated distance (in number of mutations) between each sequence and Wuhan reference sequence NC_045512.2. In another experiment we used EMBOSS cons online tool [1, 16] to calculate consensus sequence for given dataset. Then we calculated the distance between each sequence and consensus sequence obtained using EMBOSS cons. Then we calculated site frequency spectra from distances obtained in previous step (see the next subsection).

### 2.5 Site frequency spectrum and its properties

Inference from evolutionary models of DNA often exploits summary statistics of sequence data, a common one being the so-called Site Frequency Spectrum. In a sequencing experiment with a known number of sequences, we can estimate for each site at which a novel somatic mutation has arisen, the number of genomes that carry that mutation. These numbers are then grouped into sites that have the same number of copies of a mutant. Figure 1 (based on [5]; modified) gives an example with time running down the page. The genealogy of a sample of *n* = 20 cells includes 13 mutational events. We can see that mutations 4, 5, 7, 10, 11, 12, and 13 (a total of 7 mutations) are present in a single genome, mutations 1, 2, and 3 (total of 3 mutations) are present in 3 genomes, mutations 8 and 9 (a total of 2 mutations) are present in six genomes, and mutation 6 is present in 17 genomes. If we denote the number of mutations present in *k* genomes by *S_n_*(*k*), we see that in this example, *S_n_*(1) = 7, *S_n_*(3) = 3, *S_n_*(6) = 2, and *S_n_*(17) = 1, with all other *S_n_*(*j*) equal to 0. The vector (*S_n_*(1), *S_n_*(2),…, *S_n_*(*n* – 1)) is called the (observed) Site Frequency Spectrum, abbreviated to SFS. It is conventional to include only sites that are segregating in the sample, that is, those for which the mutant type and the ancestral type are both present in the sample at that site. Mutations that occur prior to the most recent common ancestor of the sampled genomes will be present in all genomes in the sample; these are not segregating and are called truncal mutations.

**Figure 1:**
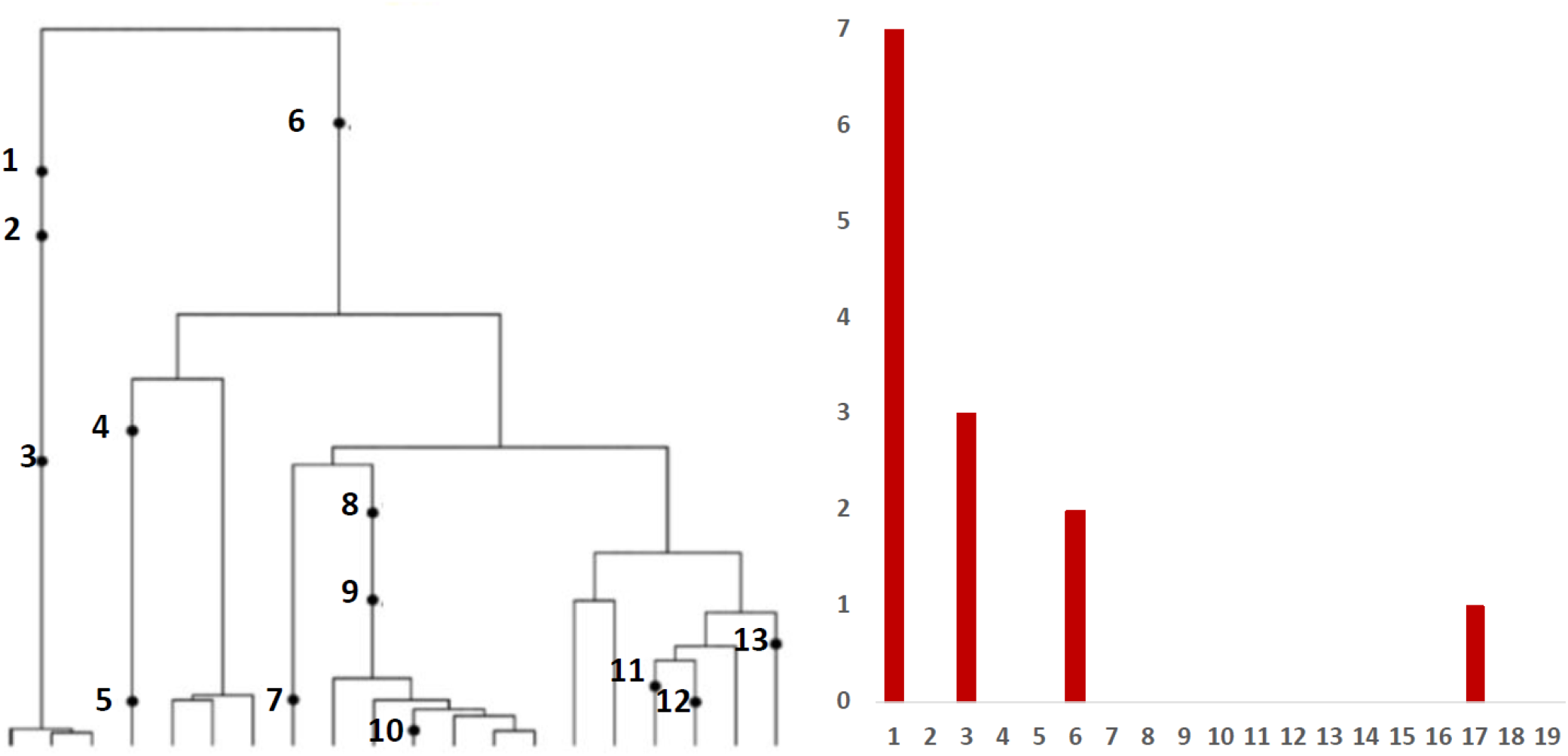
Left panel: Genealogy of a sample of *n* = 20 genomes includes 13 mutational events, denoted by black dots. Mutations 4, 5, 7, 10, 11, 12, and 13 (total of 7 mutations) are present in a single genome, mutations 1, 2, and 3 (total of 3 mutations) are present in three genomes, mutations 8 and 9 (2 mutations) are present in six genomes, and mutation 6 (1 mutation) is present in 17 genomes. Right panel: The observed site frequency spectrum, *S*_20_(1) = 7, *S*_20_(3) = 3, *S*_20_(6) = 2, and *S*_20_(17) = 1, other *S_n_*(*k*) equal to 0.

## 3 Results

### 3.1 Mutation timeline

Based on the data from processing of subsequences containing segregating sites for Alpha and Beta SARS-CoV-2 variant, we generated timelines for each of defining mutations (Fig. 2 A and B. In Fig. 3 A and B, we present cumulative plots showing changes in the dynamics of the increase in the count of individual mutations over time. We observed that although genomes containing complete set of VOC-defining mutations emerged late in 2020 (September 20, for the Alpha variant, and October 10, for the Beta variant) specific mutations emerged even in first weeks of the pandemic. This is especially true for such important mutation as D614G, classified as selectively advantageous [20]. This mutation was first detected in a genome collected in the second week of pandemic. Complete or near-complete sets of VOC-defining mutations emerged earlier in case of Alpha variant than of Beta variant.

**Figure 2:**
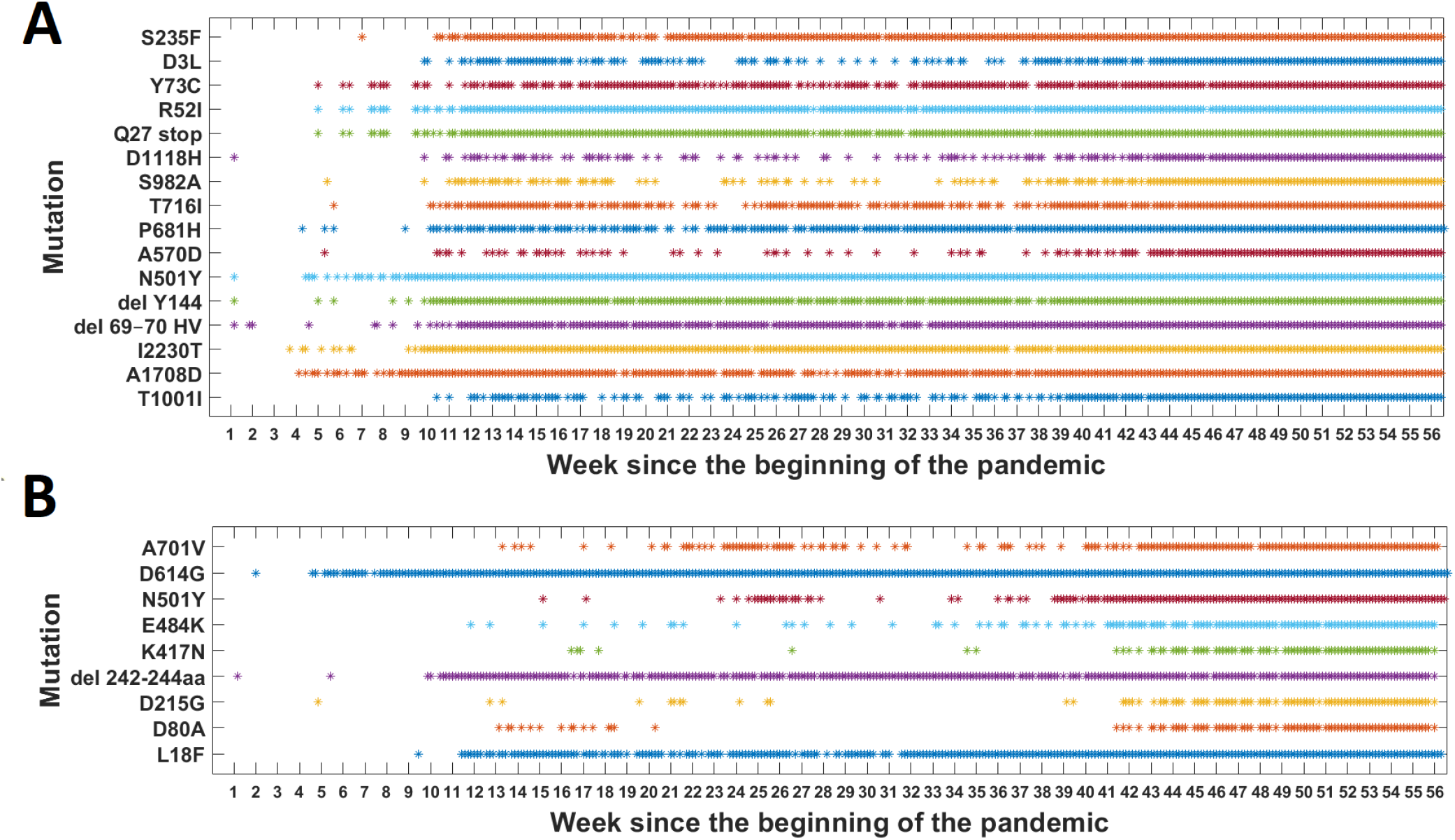
Genomes with VOC-defining mutations over time. (A) B.1.1.7 (Alpha) variant; (B) B.1.351 (Beta) variant.

**Figure 3:**
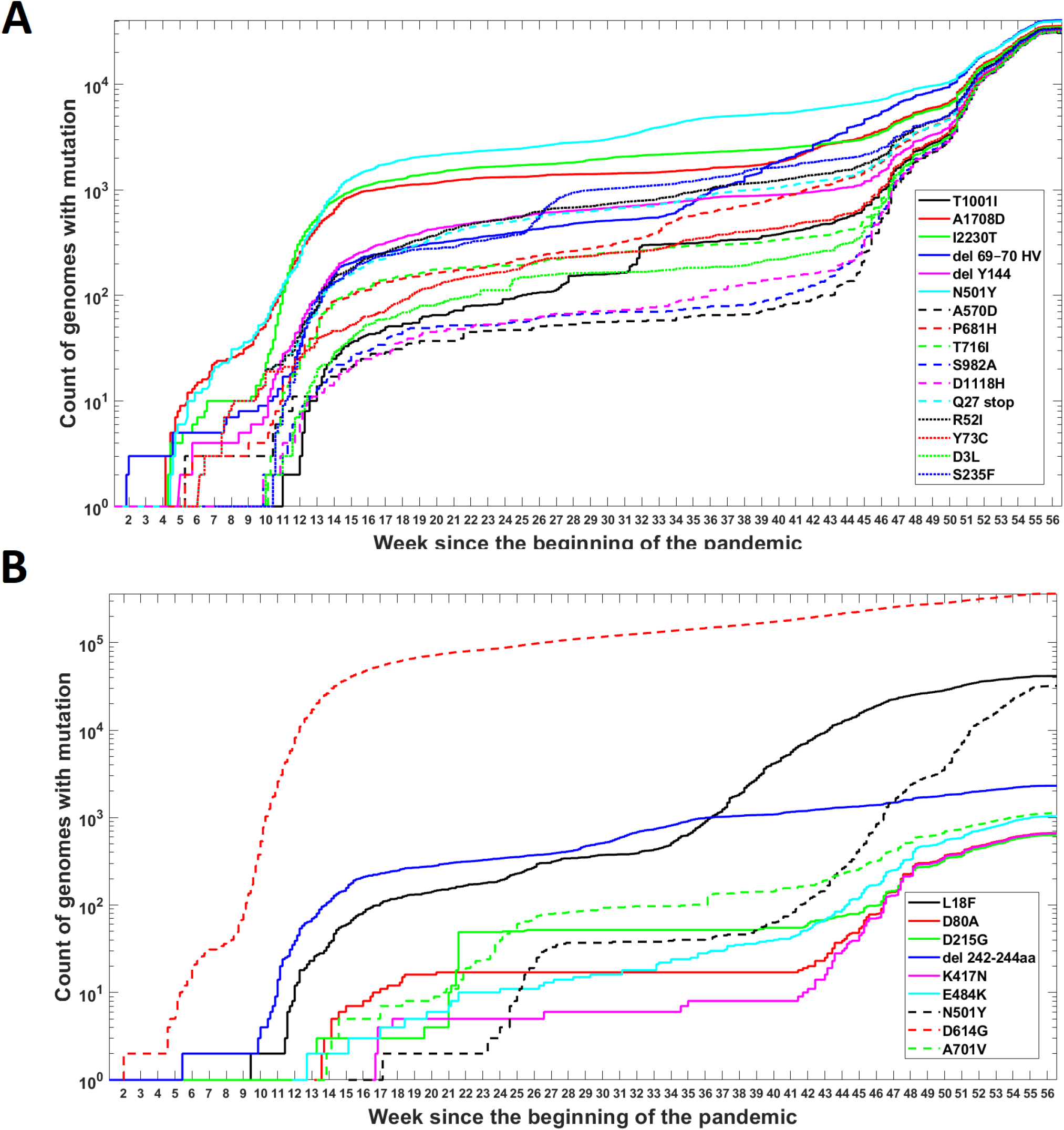
Cumulative count of genomes with VOC-defining mutations over time. (A) B.1.1.7 (Alpha) variant; (B) B.1.351 (Beta) variant.

The dynamics of increase in cumulative number of genomes with VOC-defining mutations is characterized by several humps. This is mainly caused by differences in the overall count of genomes sequenced on a given day (data were not normalized). However, for several mutations we observe a rapid increase in count which is not simply explainable by the overall number of sequenced genomes (but may be influenced by geographic factors). Examples of such mutations in the case of Alpha variant are S235F, P681H and 21765-21770 deletion. For Beta variant we observe the fastest increase in number of genomes carrying N501Y mutation.

### 3.2 Mutation frequency and combinations

For both Alpha and Beta variant we calculated how many genomes carry a given number of mutations from the VOC-defining set (Fig. 4 A and B). For both variants, we observe that there is a large number of genomes carrying only one or two from VOC-defining mutations but – especially in case of Alpha variant – there is also a lot of sequences containing the complete set (28,994 genomes for Alpha variant and 193 genomes for Beta variant). The least numerous are genomes having mutations in 4-6 out of all segregating sites.

**Figure 4:**
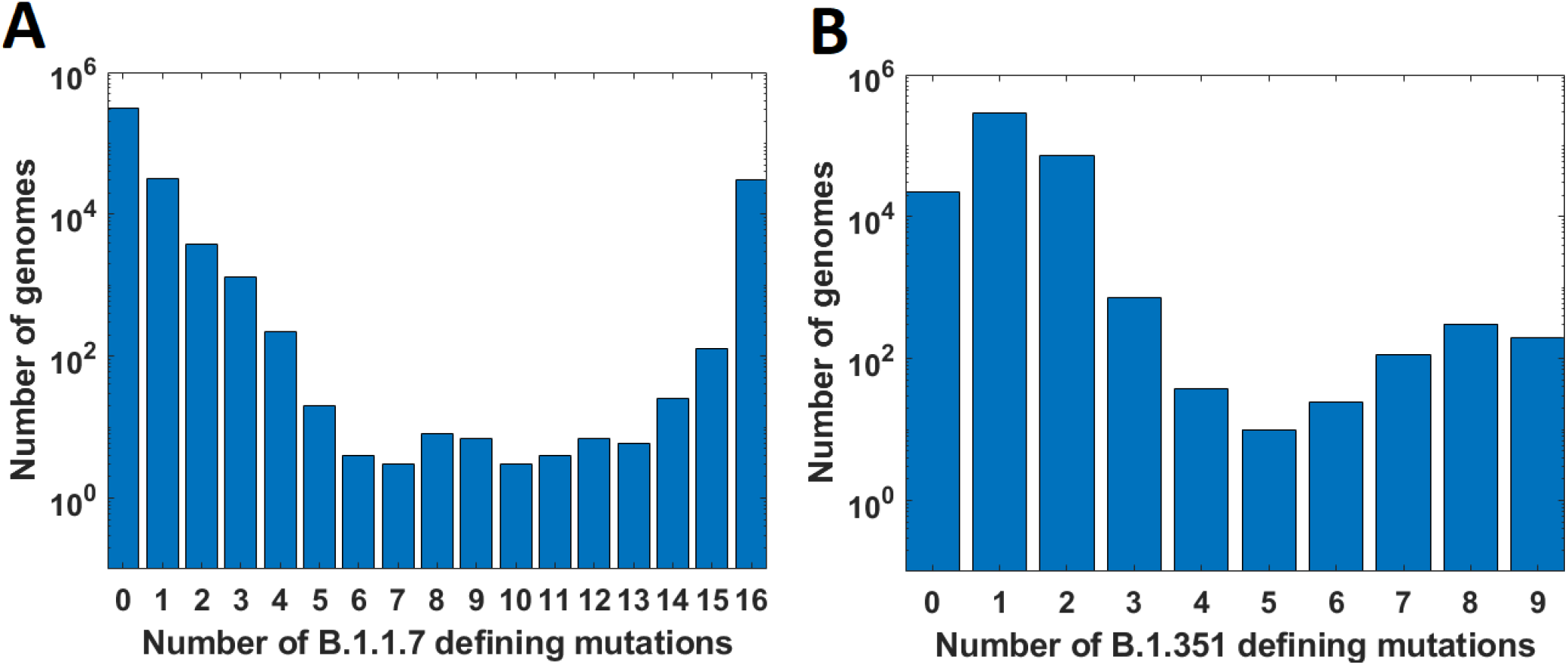
Frequency of genomes carrying given number if VOC-defining mutations. (A) B.1.1.7 (Alpha) variant; (B) B.1.351 (Beta) variant.

We calculated the number of observed unique combinations of VOC-defining mutations for both Alpha (Table 3) and Beta (Table 4) variants and compared them with expected number of combinations for given mutation count. Results presented in Fig. 5 A and B and in Tables 3 and 4 clearly show that obtained results depart from expectations.

**Figure 5:**
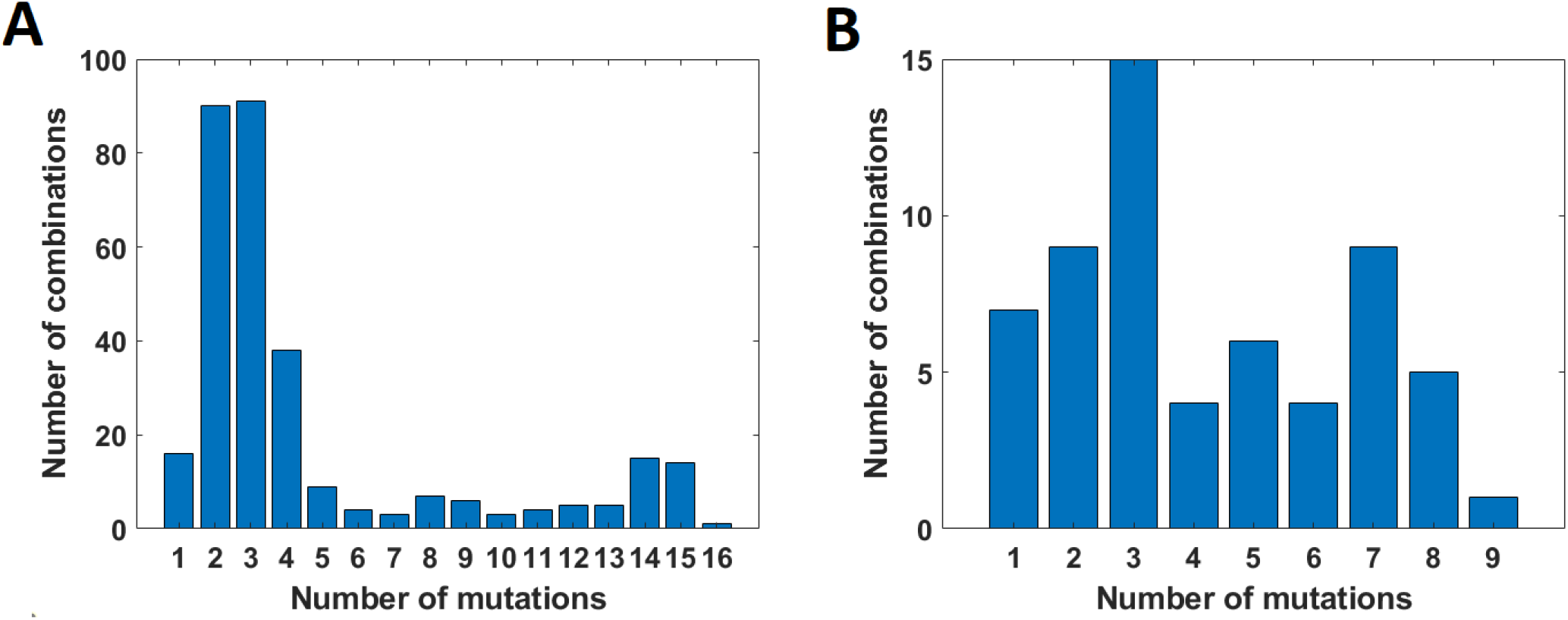
Number of unique combinations of VOC-defining mutations for given mutation count. (A) B.1.1.7 (Alpha) variant; (B) B.1.351 (Beta) variant.

**Table 3:** Number of unique combinations of VOC Alpha-defining mutations for given mutation count, out of a total of 16.

| **Number of mutations** | **Expected** | **Observed** |
| --- | --- | --- |
| 1 | 16 | 16 |
| 2 | 120 | 90 |
| 3 | 560 | 91 |
| 4 | 1820 | 38 |
| 5 | 4368 | 9 |
| 6 | 8008 | 4 |
| 7 | 11440 | 3 |
| 8 | 12870 | 7 |
| 9 | 11440 | 6 |
| 10 | 8008 | 3 |
| 11 | 4368 | 4 |
| 12 | 1820 | 5 |
| 13 | 560 | 5 |
| 14 | 120 | 15 |
| 15 | 16 | 14 |
| 16 | 1 | 1 |

**Table 4:** Number of unique combinations of VOC Beta-defining mutations for given mutation count, out of a total of 9.

| **Number of mutations** | **Expected** | **Observed** |
| --- | --- | --- |
| 1 | 9 | 7 |
| 2 | 36 | 9 |
| 3 | 84 | 15 |
| 4 | 126 | 4 |
| 5 | 126 | 6 |
| 6 | 84 | 4 |
| 7 | 36 | 9 |
| 8 | 9 | 5 |
| 9 | 1 | 1 |

We investigated when combinations of given number of mutations first emerge in time and what is the dynamics of increase in number of unique combinations over time (Fig. 6). We observe that genomes carrying combinations of higher number of mutations (even full set) emerge earlier than genomes carrying only some of them (e.g. combinations of 5 or 6 mutations).

**Figure 6:**
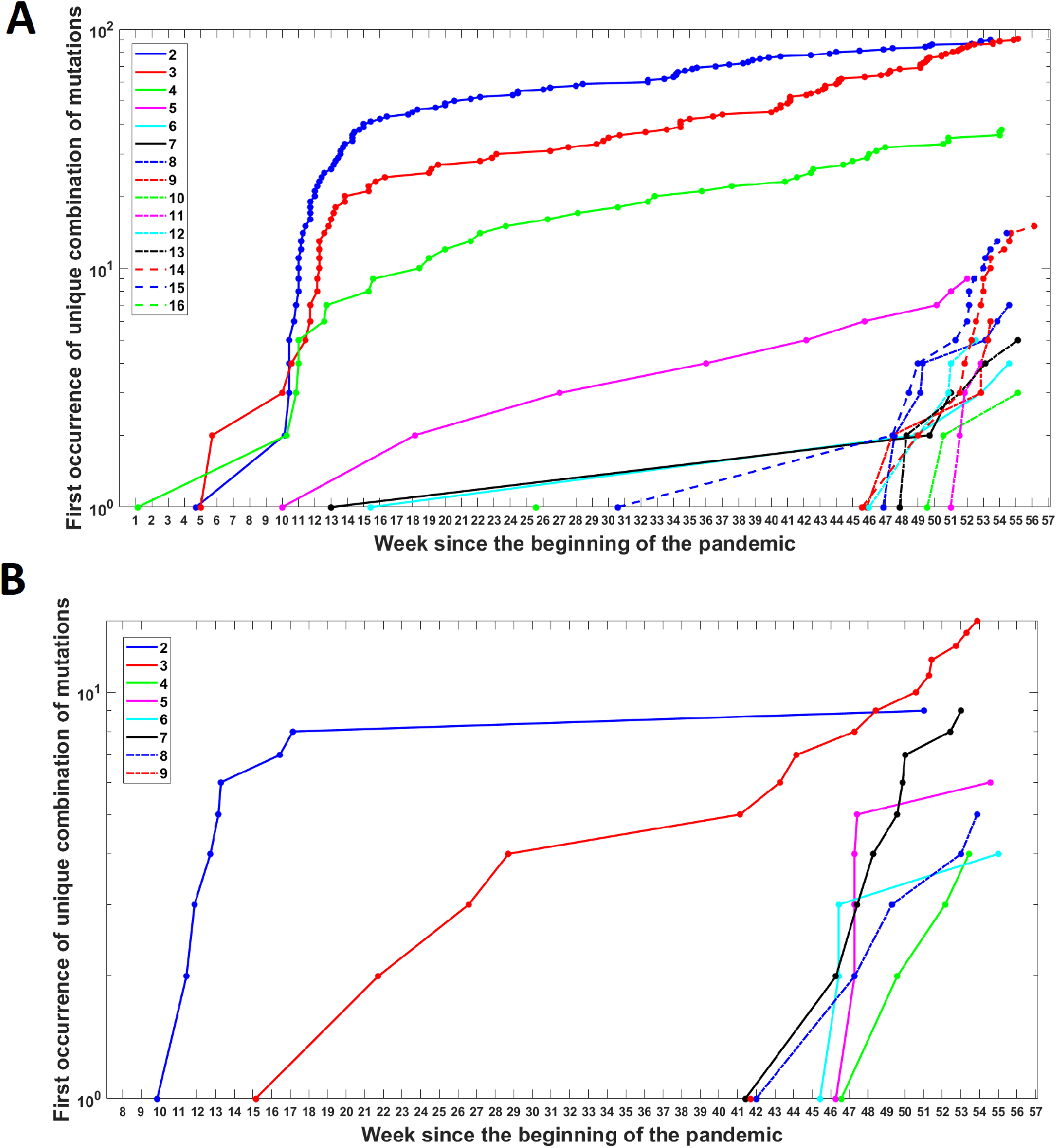
Cumulative count of unique combinations of given number of mutations over time. (A) B.1.1.7 (Alpha) variant; (B) B.1.351 (Beta) variant.

### 3.3 Studies on the site frequency spectrum

The results of analysis of the genomes sequenced in week 45, 46 and 49 for Alpha variant and all genomes for Beta variant are presented in the form of log-log cumulative tails (Figs 7–8 and S2–S3). We observe that the slope of cumulative tails differs between sample with Alpha genomes and the sample with remaining ones. In case of Beta variant we do not observe such significant effect.

**Figure 7:**
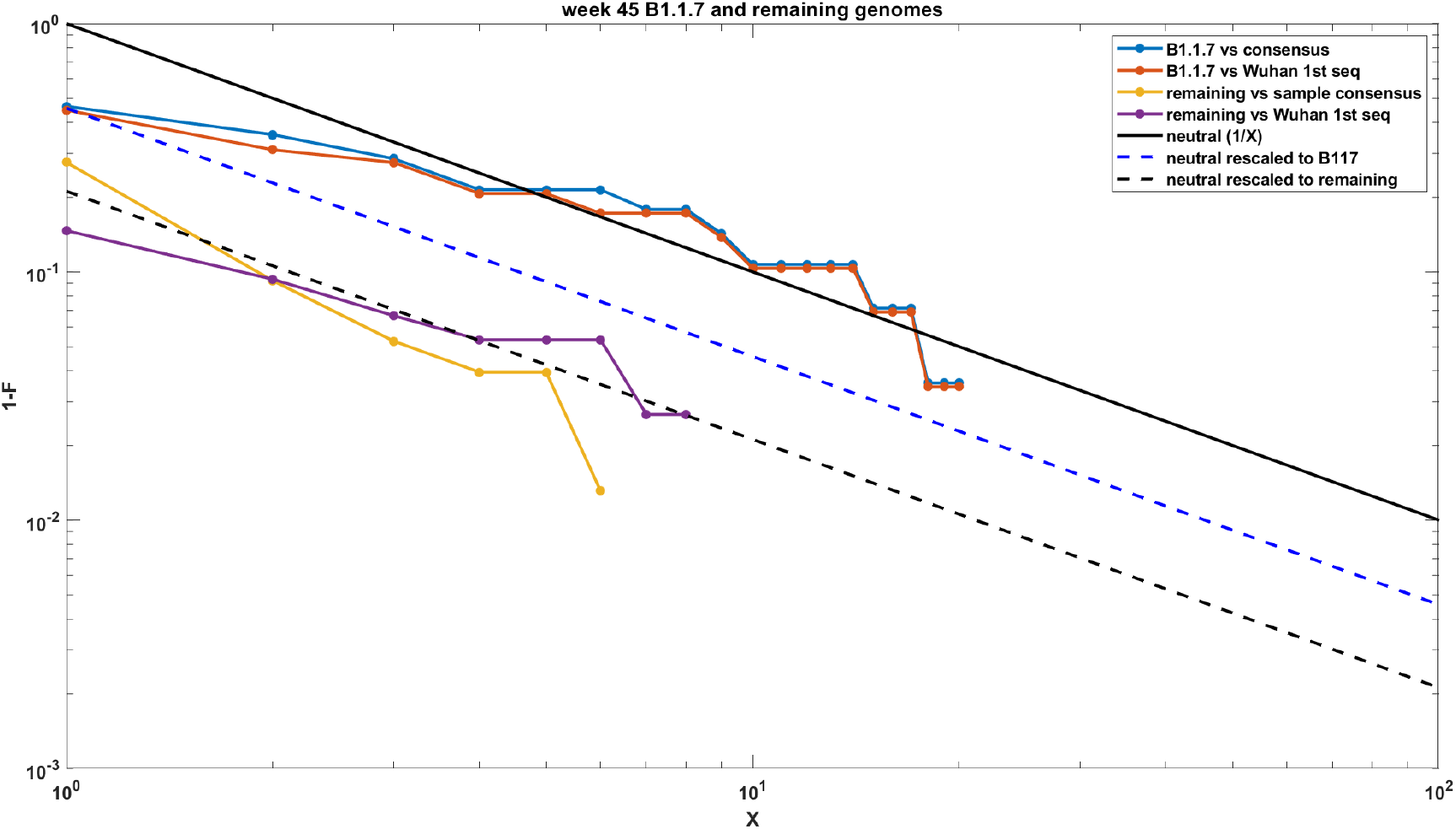
Comparison of SFS cumulative tails calculated from mutations present in B.1.1.7 genomes with SFS cumulative tails calculated from mutations present in equivalent number of remaining genomes in week 45.

**Figure 8:**
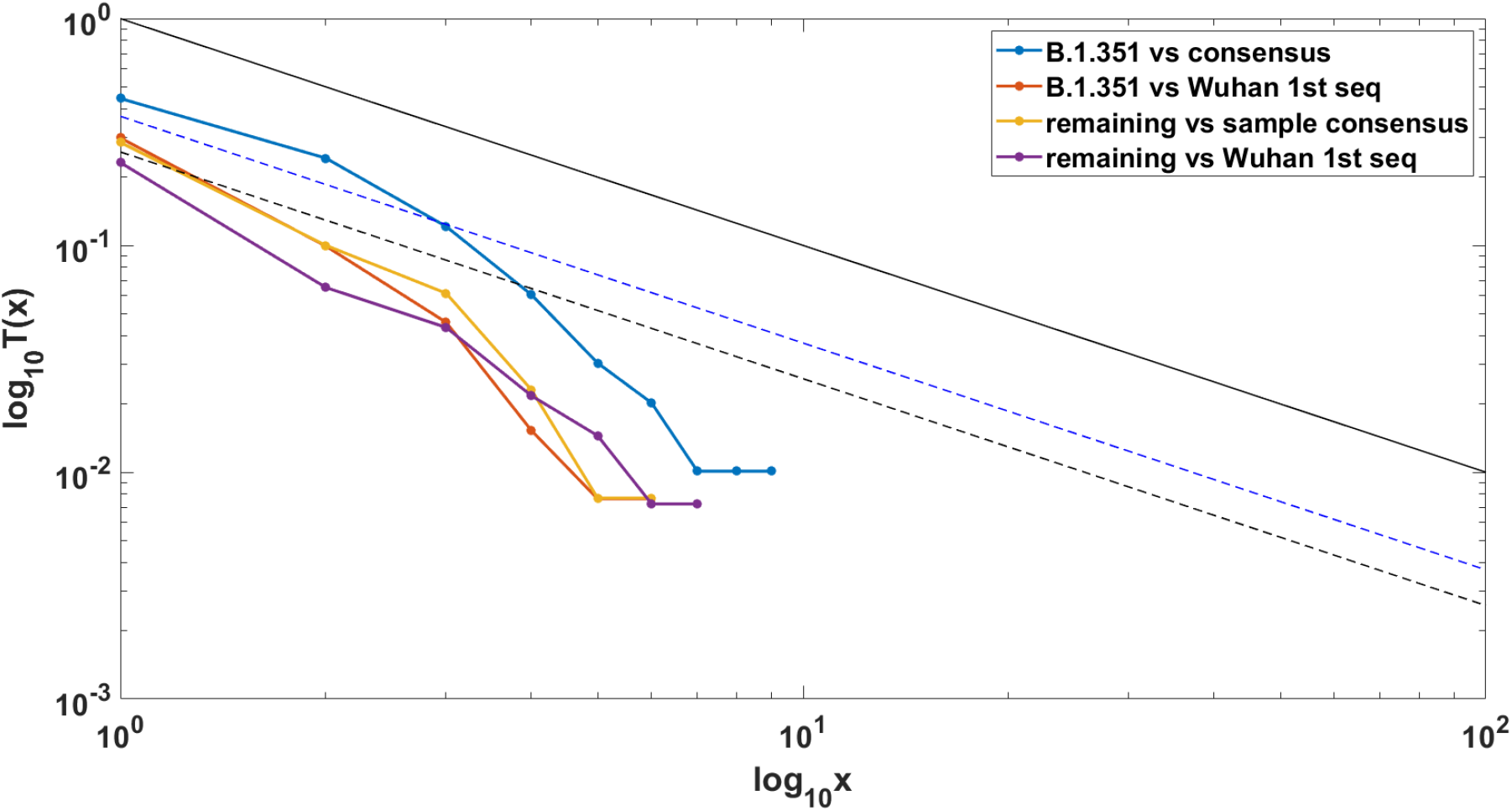
Comparison of SFS cumulative tails calculated from mutations present in B.1.351 genomes with SFS cumulative tails calculated from mutations present in equivalent number of remaining genomes.

In the case of exponential population growth, Durrett [6] provided an approximate large sample and large population expression, which leads to the conclusion that, assuming neutral evolution, the SFS cumulative tail in the log-log scale should be approximated by a straight line with coefficient −1 (marked in Figs 7–8 and S2-S3).

Analysis of the obtained results shows that cumulative SFS tails calculated based on non-Alpha genomes data can be approximated by straight line with coefficient −1 characteristic for neutral evolution. On the contrary, the slope of cumulative SFS tails obtained for Alpha variant genomes indicates the presence of selective pressure on the evolution of these genomes.

## 4 Discussion

In this study we analysed SARS-CoV-2 genomes to see how the individual mutations that define the Alpha and Beta variants were appearing over time. Our analyses showed that these mutations did not arose gradually, but rather co-evolved rapidly leading to the emergence of the full VOC strain. We do not observe transient states which would be expected under neutral evolution. These results seem to indicate that segregating sites in Alpha and Beta variants evolved under strong positive selection. Another possible explanation might be recombination event between viruses carrying subsets of VOC-defining mutations. Research has shown that such phenomenon is common in bat coronaviruses [13] and might be indeed affecting also the evolution of SARS-CoV-2 [11]. Observed mutation patterns may be also due to mutation hotspots, which were detected in the region encoding the Spike protein [14].

In addition to the factors described above, we cannot rule out the possibility that genomes carrying subsets of VOC-defining mutations avoided collection and sequencing. In the data gathered by GISAID we can clearly see temporal differences in the number of sequenced genomes (as shown in Fig. S1) but more importantly most of collected genomes come from Europe and United States. The under representation of sequences from other parts of the world could possibly be the cause why genomes containing subsets of mutations have been overlooked.

We carried out additional analysis of the early evolution of the B1.1.7 VOC, in the week 45 of the epidemic, when only 115 samples of the variant were present and its abundance was still increasing roughly exponentially. We used the model developed in [5]. The model assumes that at some time labeled *t*_0_ = 0, strain of viruses, such as the VOC B.1.7.7 (clone 0) arises, grows deterministically in size at rate *r*_0_, these cells acquiring mutations at the rate *θ*_0_ per time unit per genome. At time *t*_1_ > 0, a subclone (clone 1) arises, which differs from the original clone with respect to growth rate (now equal to *r*_1_ > *r*(0)) and mutation rate (now equal to *θ*_1_). We call this the “selective event”. The new clone arises on the background of a haplotype already harboring *K* mutations. Finally, at *t*_2_ > *t*_1_ > 0, a sample of *n* variant’s RNA genomes is sequenced. Without getting into details, as explained in [5], the emerging substrain leaves a signature (“bulge”) on the SFS cumulative tail *T*(*x*), the characteristics of which can be estimated from equations in [5]. Figure 9 illustrates the fit. The conclusion is that we observe a substrain of B.1.1.7, which has estimated *K* = 25 additional mutations and which constitutes fraction of *p* = 0.13 of the B.1.1.7 genome. These conclusions, if confirmed, suggest a high diversification within the B.1.1.7, occurring at the time of its emergence.

**Figure 9:**
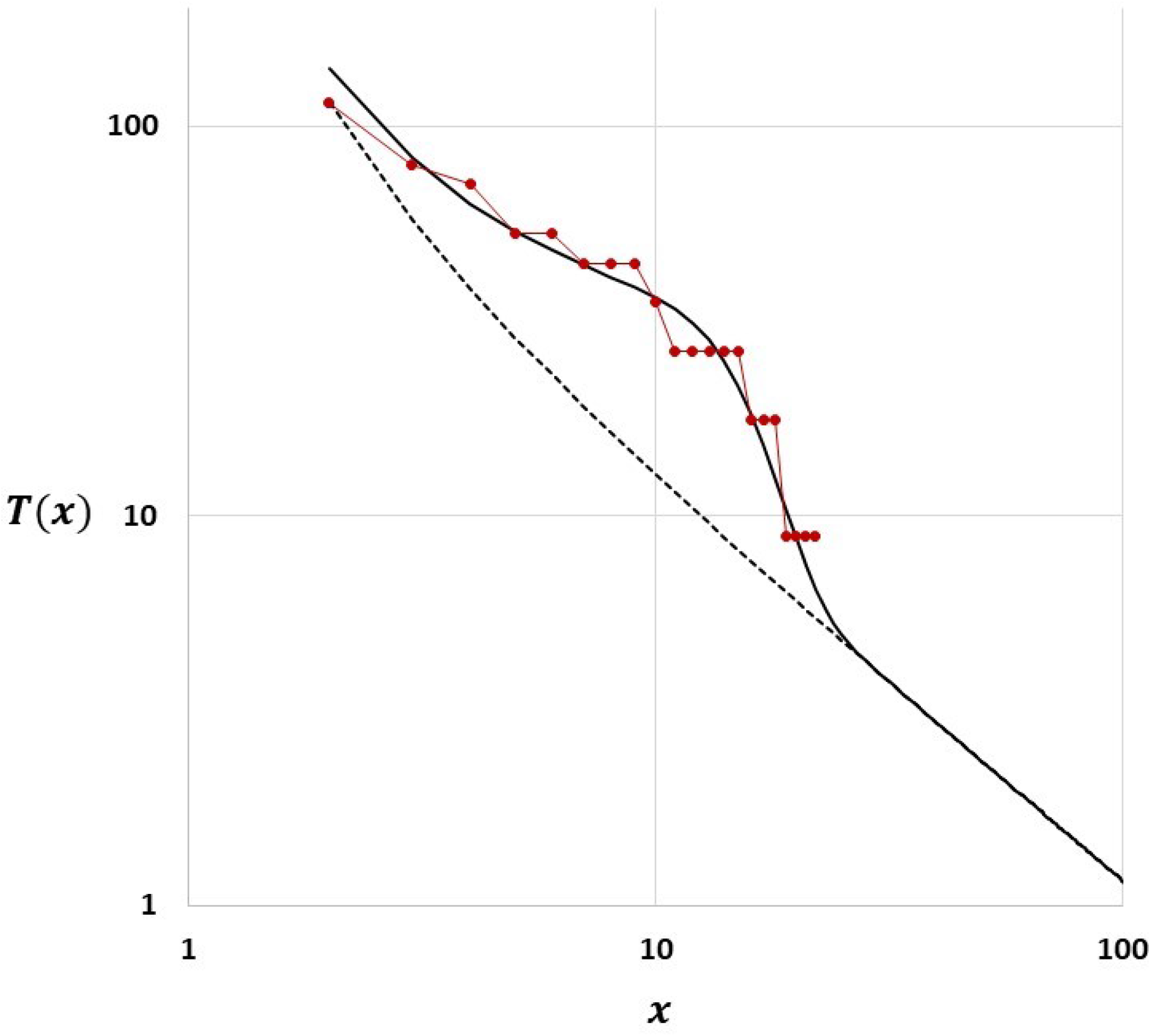
Comparison of SFS cumulative tail *T*(*x*) calculated from mutations present in B.1.1.7 genomes in week 45 (red bulets, linked with red lines for visual convenience) with SFS cumulative tail calculated from the model in [5] (continuous black line). The bulge-shaped line is the signature of a competitive subclone emerging. Dashed black slightly convex line is the reference line that the model predicts if no competitive subclone arises.

## Author’s contribution

MKi suggested the problem, designed and supervised the research. MKu designed algorithms to collect weekly statistics of viral genomes. MKu performed the analyses and visualized the results. MKi and MKu prepared the manuscript. All authors reviewed and approved the final version.

## Additional Information

### Data availability

All relevant data are included within the manuscript and the Supporting Information files.

## Supplementary Information

### Accession numbers

NC_045512.2 - Wuhan reference sequence

EPI_ISL_601443 - the first Alpha variant genome collected

EPI_ISL_712073 - the first Beta variant genome collected

**Figure S1:**
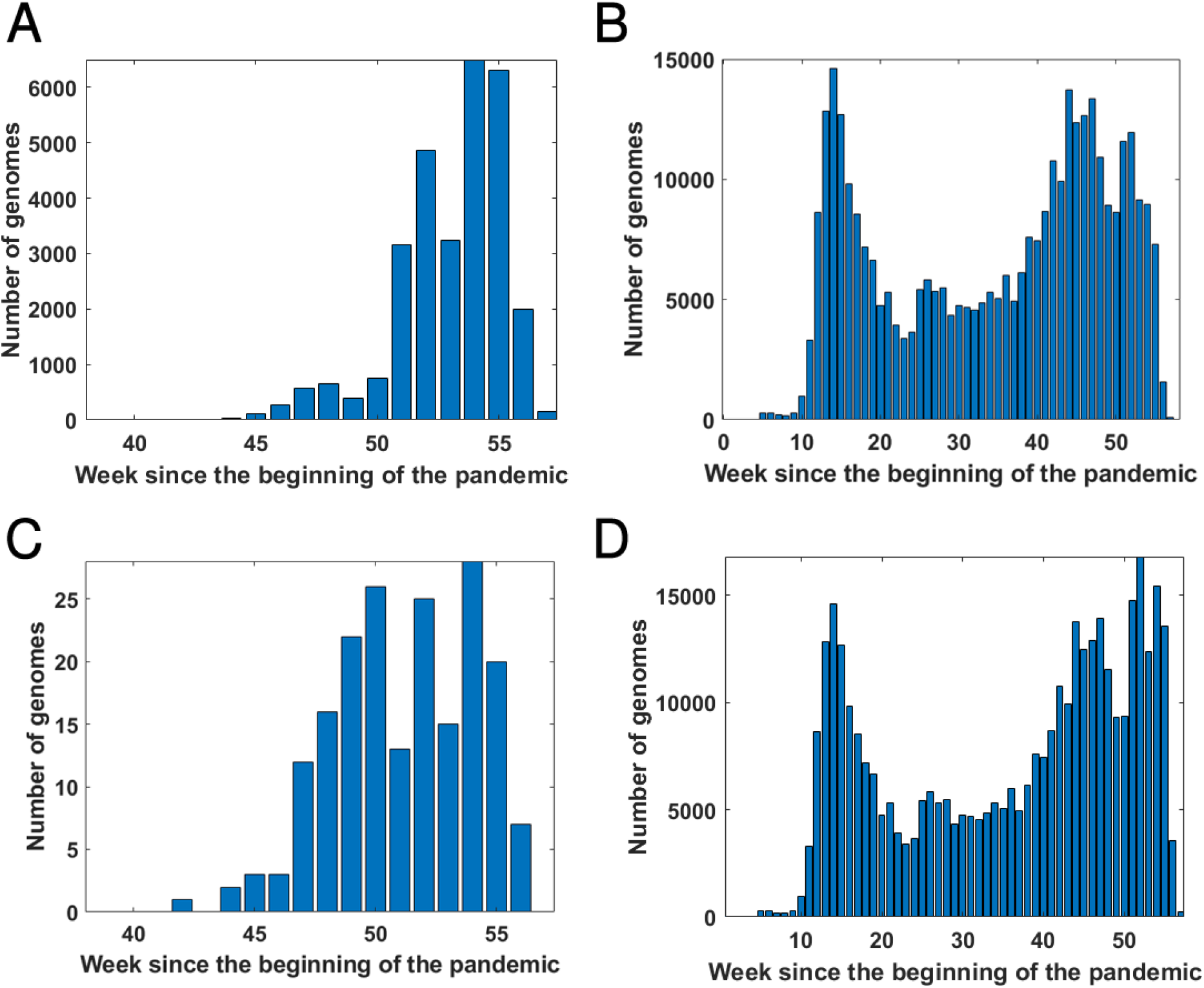
Count of VOC and remaining genomes in given weeks of the pandemic. (A) B.1.1.7 genomes; (B) Genomes other than B.1.1.7; (C) B.1.351 genomes; (D) Genomes other than B.1.351.

**Figure S2:**
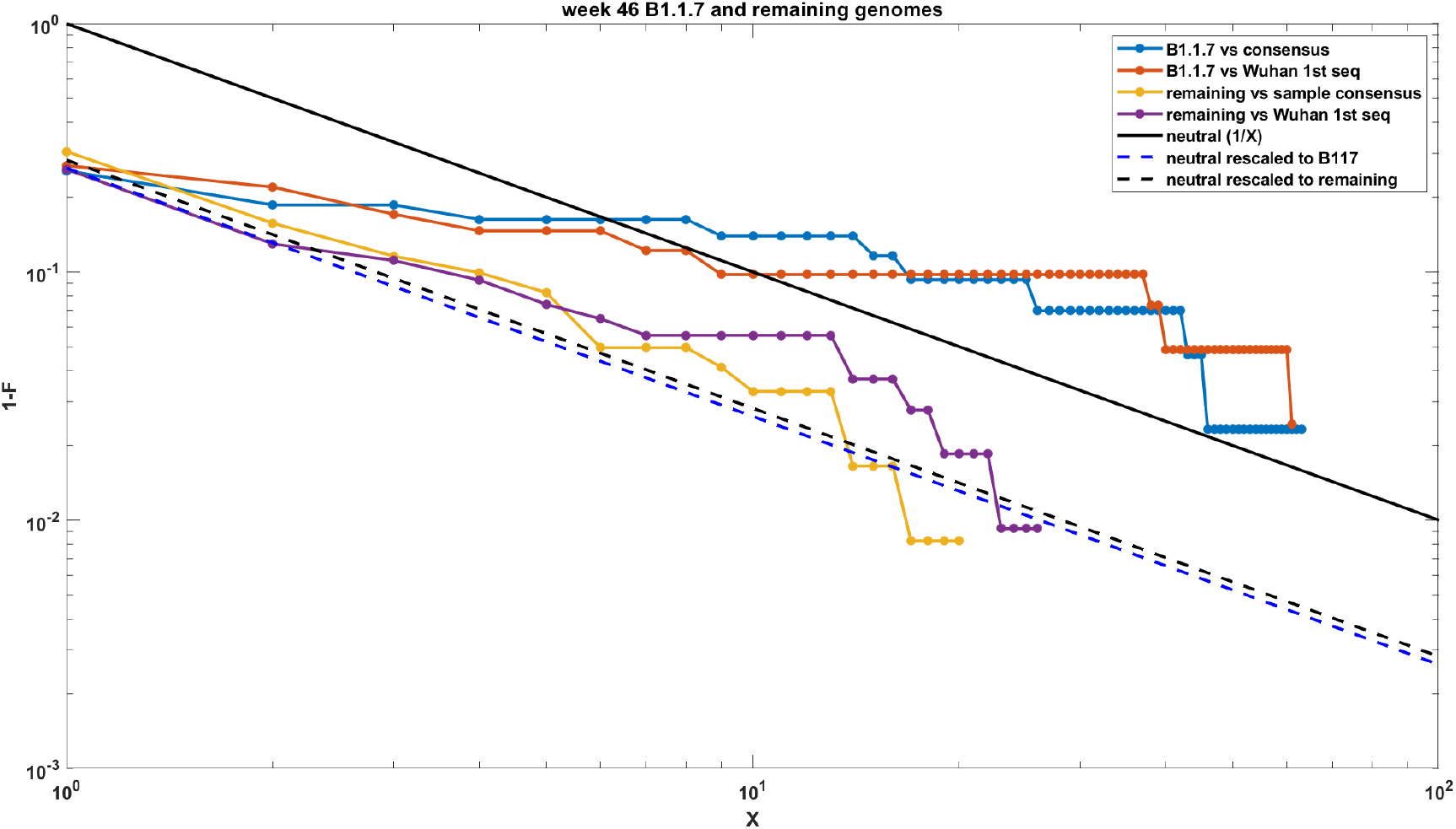
Comparison of SFS cumulative tails calculated from mutations present in B.1.1.7 genomes with SFS cumulative tails calculated from mutations present in equivalent number of remaining genomes in week 46.

**Figure S3:**
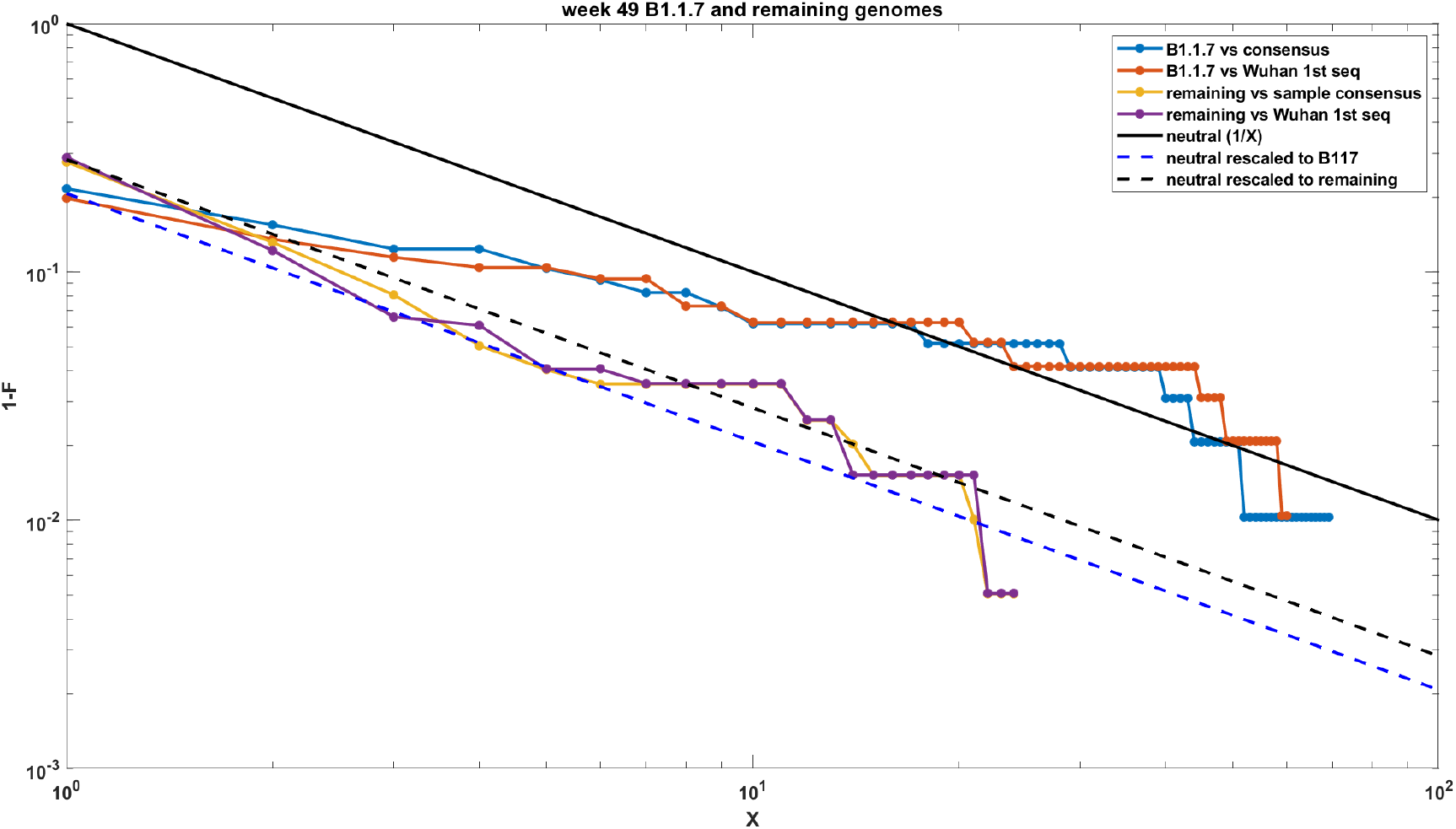
Comparison of SFS cumulative tails calculated from mutations present in B.1.1.7 genomes with SFS cumulative tails calculated from mutations present in equivalent number of remaining genomes in week 49.

## References

[1] EMBOSS: The European Molecular Biology Open Software Suite. https://www.bioinformatics.nl/cgi-bin/emboss/cons, (accessed January 29, 2022).

[2] GISAID database. https://www.gisaid.org/, (accessed January 29, 2022).

[3] Severe acute respiratory syndrome coronavirus 2 isolate Wuhan-Hu-1, complete genome. NCBI Reference Sequence: NC_045512.2. https://www.ncbi.nlm.nih.gov/nuccore/1798174254, (accessed January 29, 2022).

[4] Nicholas G Davies, Sam Abbott, Rosanna C Barnard, Christopher I Jarvis, Adam J Kucharski, James D Munday, Carl AB Pearson, Timothy W Russell, Damien C Tully, Alex D Washburne, et al. Estimated transmissibility and impact of sars-cov-2 lineage b. 1.1. 7 in england. Science, 372(6538), 2021.

[5] Khanh N Dinh, Roman Jaksik, Marek Kimmel, Amaury Lambert, Simon Tavaré, et al. Statistical inference for the evolutionary history of cancer genomes. Statistical Science, 35(1):129–144, 2020.

[6] Rick Durrett. Population genetics of neutral mutations in exponentially growing cancer cell populations. The annals of applied probability: an official journal of the Institute of Mathematical Statistics, 23(1):230, 2013.

[7] Frederic Grabowski, Grzegorz Preibisch, Stanisław Giziński, Marek Kochańczyk, and Tomasz Lipniacki. Sars-cov-2 variant of concern 202012/01 has about twofold replicative advantage and acquires concerning mutations. Viruses, 13(3):392, 2021.

[8] William T Harvey, Alessandro M Carabelli, Ben Jackson, Ravindra K Gupta, Emma C Thomson, Ewan M Harrison, Catherine Ludden, Richard Reeve, Andrew Rambaut, Sharon J Peacock, et al. Sars-cov-2 variants, spike mutations and immune escape. Nature Reviews Microbiology, 19(7):409–424, 2021.

[9] Kazutaka Katoh, Kazuharu Misawa, Kei-ichi Kuma, and Takashi Miyata. Mafft: a novel method for rapid multiple sequence alignment based on fast fourier transform. Nucleic acids research, 30(14):3059–3066, 2002.

[10] Kazutaka Katoh and Daron M Standley. Mafft multiple sequence alignment software version 7: improvements in performance and usability. Molecular biology and evolution, 30(4):772–780, 2013.

[11] Xiaojun Li, Elena E Giorgi, Manukumar Honnayakanahalli Marichannegowda, Brian Foley, Chuan Xiao, Xiang-Peng Kong, Yue Chen, S Gnanakaran, Bette Korber, and Feng Gao. Emergence of sars-cov-2 through recombination and strong purifying selection. Science Advances, 6(27):eabb9153, 2020.

[12] Roujian Lu, Xiang Zhao, Juan Li, Peihua Niu, Bo Yang, Honglong Wu, Wenling Wang, Hao Song, Baoying Huang, Na Zhu, et al. Genomic characterisation and epidemiology of 2019 novel coronavirus: implications for virus origins and receptor binding. The lancet, 395(10224):565–574, 2020.

[13] Oscar A MacLean, Spyros Lytras, Steven Weaver, Joshua B Singer, Maciej F Boni, Philippe Lemey, Sergei L Kosakovsky Pond, and David L Robertson. Natural selection in the evolution of sars-cov-2 in bats created a generalist virus and highly capable human pathogen. PLoS biology, 19(3):e3001115, 2021.

[14] Baishali Mullick, Rishikesh Magar, Aastha Jhunjhunwala, and Amir Barati Farimani. Understanding mutation hotspots for the sars-cov-2 spike protein using shannon entropy and k-means clustering. Computers in biology and medicine, 138:104915, 2021.

[15] Andrew Rambaut, Nick Loman, Oliver Pybus, Wendy Barclay, Jeff Barrett, Alesandro Carabelli, Tom Connor, Tom Peacock, David L Robertson, Erik Volz, et al. Preliminary genomic characterisation of an emergent sars-cov-2 lineage in the uk defined by a novel set of spike mutations. Genom. Epidemiol, pages 1–5, 2020.

[16] Peter Rice, Ian Longden, and Alan Bleasby. Emboss: the european molecular biology open software suite. Trends in genetics, 16(6):276–277, 2000.

[17] Yuelong Shu and John McCauley. Gisaid: Global initiative on sharing all influenza data–from vision to reality. Eurosurveillance, 22(13):30494, 2017.

[18] Shuo Su, Gary Wong, Weifeng Shi, Jun Liu, Alexander CK Lai, Jiyong Zhou, Wenjun Liu, Yuhai Bi, and George F Gao. Epidemiology, genetic recombination, and pathogenesis of coronaviruses. Trends in microbiology, 24(6):490–502, 2016.

[19] Houriiyah Tegally, Eduan Wilkinson, Marta Giovanetti, Arash Iranzadeh, Vagner Fonseca, Jennifer Giandhari, Deelan Doolabh, Sureshnee Pillay, Emmanuel James San, Nokukhanya Msomi, et al. Detection of a sars-cov-2 variant of concern in south africa. Nature, 592(7854):438–443, 2021.

[20] Erik Volz, Verity Hill, John T McCrone, Anna Price, David Jorgensen, Aine O’Toole, Joel Southgate, Robert Johnson, Ben Jackson, Fabricia F Nascimento, et al. Evaluating the effects of sars-cov-2 spike mutation d614g on transmissibility and pathogenicity. Cell, 184(1):64–75, 2021.

